# Application of monolayer graphene to cryo-electron microscopy grids for high-resolution structure determination

**DOI:** 10.1101/2023.07.28.550908

**Authors:** Andrew V. Grassetti, Mira B. May, Joseph H. Davis

**Affiliations:** Department of Biology, Massachusetts Institute of Technology, Cambridge, MA 02139; Program in Computational and Systems Biology, Massachusetts Institute of Technology, Cambridge, MA 02139

## Abstract

In cryogenic electron microscopy (cryo-EM), purified macromolecules are typically applied to a grid bearing a holey carbon foil, blotted to remove excess liquid and rapidly frozen in a roughly 20-100 nm thick layer of vitreous ice that is suspended across roughly 1 μm-wide foil holes. The resulting sample is then imaged using cryogenic transmission electron microscopy and, after substantial image processing, near-atomic resolution structures can be determined. Despite cryo-EM’s widespread adoption, sample preparation remains a severe bottleneck in cryo-EM workflows, with users often encountering challenges related to samples behaving poorly in the suspended vitreous ice. Recently, methods have been developed to modify cryo-EM grids with a single continuous layer of graphene, which acts as a support surface that often increases particle density in the imaged area and can reduce interactions between particles and the air-water interface. Here, we provide detailed protocols for the application of graphene to cryo-EM grids, and for rapidly assessing the relative hydrophilicity of the resulting grids. Additionally, we describe an EM-based method to confirm the presence of graphene by visualizing its characteristic diffraction pattern. Finally, we demonstrate the utility of these graphene supports by rapidly reconstructing a 2.7 Å resolution density map of an exemplar Cas9 complex using a highly pure sample at a relatively low concentration.

**SUMMARY:** The application of support layers, such as graphene, to cryo-electron microscopy grids can increase the density of particles imaged, limit particle interactions with the air-water interface, reduce the extent of beam-induced motion, and, in some instances, improve the distribution of particle orientations. This paper describes a robust protocol for coating cryo-EM grids with a monolayer of graphene for improved cryo-sample preparation.

## INTRODUCTION

Single particle cryo-electron microscopy (cryo-EM) has evolved into a widely used method for visualizing biological macromolecules^1^. Fueled by advances in direct electron detection^2-4^, data acquisition^5^, and image processing algorithms^6-10^, cryo-EM is now capable of producing near-atomic resolution 3D structures of a fast growing number of macromolecules^11^. Moreover, by leveraging the single-molecule nature of the approach, users can now determine multiple structures from a single sample^12-15^, highlighting the promise of using these data to understand heterogeneous structural ensembles^16,17^. Despite this progress, bottlenecks in cryo-specimen grid preparation persist.

For structural characterization by cryo-EM, biological samples should be well-dispersed in aqueous solution and then must be flash-frozen through a process called vitrification^18,19^. The goal is to capture particles in a uniformly thin layer of vitrified ice suspended across regularly-spaced holes that are typically cut into a layer of amorphous carbon. This patterned amorphous carbon foil is supported by a TEM grid bearing a mesh of copper or gold support bars. In standard workflows, grids are rendered hydrophilic using a glow-discharge plasma treatment prior to the application of sample. Excess liquid is blotted with filter paper, allowing the protein solution to form a thin liquid film across the holes that can be readily vitrified during plunge-freezing. Common challenges include particle localization to the air-water interface (AWI) and subsequent denaturation^20-22^ or adoption of preferred orientations^23-25^, particle adherence to the carbon foil rather than migrating into the holes, and clustering and aggregation of the particles within the holes^26^. Nonuniform ice thickness is another concern; thick ice can result in higher levels of background noise in the micrographs due to increased electron scattering, whereas extremely thin ice can exclude larger particles^27^.

To address these challenges, a variety of thin support films have been used to coat grid surfaces, allowing particles to rest on these supports and, ideally, avoid interactions with the air-water interface. Graphene supports have shown great promise, in part due to their high mechanical strength coupled with their minimal scattering cross-section, which reduces the background signal added by the support layer^28^. In addition to its minimal contribution to background noise, graphene also exhibits remarkable electrical and thermal conductivity^29^. Graphene and graphene oxide coated grids have been shown to yield higher particle density, more uniform particle distribution^30^, and reduced localization to the AWI^22^. In addition, graphene provides a support surface that can be further modified to: 1) tune the physiochemical properties of the grid surface through functionalization^31-33^; or 2) couple linking agents that facilitate affinity purification of proteins of interest^34-36^.

In this article, we have modified an existing procedure for coating cryo-EM grids with a single uniform layer of graphene^30^. Our modifications aim to minimize grid handling throughout the protocol, with the goal of increasing yield and reproducibility. Additionally, we discuss our approach to evaluate the efficacy of various UV/ozone treatments in rendering grids hydrophilic prior to plunging. This step in cryo-EM sample preparation using graphene-coated grids is critical, and we have found our straightforward method to quantify the relative hydrophilicity of the resulting grids to be useful. Using this protocol, we demonstrate the utility of employing graphene-coated grids for structure determination by generating a high-resolution 3D reconstruction of catalytically inactive *S. pyogenes* Cas9 in complex with guide RNA and target DNA.

## PROTOCOL

### 1 Fabrication of graphene-coated cryo-EM grids

#### 1.1 Prepare graphene etching solution

- Dissolve 4.6 g ammonium persulfate (APS) in 20 mL molecular grade water in a 50 mL beaker for a 1 M solution and cover with aluminum foil.
- Allow APS to completely dissolve while proceeding to step 1.2.

#### 1.2 Prepare a section of CVD Graphene for methyl methacrylate (MMA) coating

- Carefully cut a square section of CVD graphene.
- Transfer the square to a coverslip within a clean petri dish and cover during transport to the spin coater. **NOTE:** An 18 mm x 18 mm square should yield 25-36 graphene-coated grids.

#### 1.3 Coating CVD graphene with MMA

- Initialize the spin coater by pressing the red power switch on the bottom left of the instrument.
- Turn on the black pump power switch next to the plug port.
- Set the spin coater settings to a 60-second high speed spin at 2,500 rpm.
- Carefully place the CVD graphene on the appropriately sized chuck. **NOTE:** Ideally, the CVD graphene will extend 1-2 mm over the edge of the selected chuck to prevent aspiration of MMA into the vacuum system.
- Press the **Take/Absorb** button to engage the vacuum pump and affix the CVD graphene to the chuck.
- Apply MMA to the center of CVD graphene square, close lid and press **Start/ Stop** button immediately. **NOTE:** 40 μL of MMA is appropriate for an 18 mm x 18 mm square of CVD graphene.
- Once spinning has stopped, press the **Take/Absorb** button to disengage the vacuum pump and release the MMA coated CVD graphene.
- Carefully retrieve the MMA coated CVD graphene with tweezers.
- Invert the CVD graphene such that the MMA coated side is facing down and place it back on the glass coverslip.

#### 1.4 Plasma etching of the graphene back-side

- Transfer CVD graphene on the coverslip into the PELCO easiGlow and glow discharge for 30 seconds at 25 mA using flat-tip tweezers.
- Return the CVD graphene on the coverslip to the petri dish and cover during transport to the copper etching area. **NOTE:** MMA coating will protect the topside graphene layer from plasma etching.

#### 1.5 Cut grid-sized squares and dissolve copper substrate

- Use two pairs of tweezers to support the CVD graphene square. **CRITICAL:** Take note of the orientation of the CVD graphene square, it should be MMA side up while attached to the tweezers.
- Cut CVD graphene into approximately 3 mm x 3 mm squares. **NOTE:** Using two sets of reverse-action forceps for this step allows one to hold the CVD graphene square right-side-up and anchor it in position, and also to hold the edge of a 3 mm x 3 mm square after cutting it away from the rest of the square.
- Carefully float each 3 mm x 3 mm square in the APS solution.
- Make contact with the surface of the APS solution prior to releasing each square. **NOTE:** Tilting the beaker such that you can place the square into the solution at a shallow angle helps ensure that the square does not sink.
- Cover beaker with aluminum foil and incubate overnight at 25°C. **NOTE:** If you do not have a glass coverslip cut to dimensions 12 mm x 50 mm, use a glass cutter and a straightedge to cut a 24 mm x 50 mm coverslip in half.

#### 1.6 Remove MMA/graphene films from APS and incubate in water

- Use a glass coverslip with dimensions 12 mm x 50 mm and extract the MMA/ graphene squares from APS by gently plunging the coverslip vertically into the APS and then moving the coverslip laterally such that it abuts a floating MMA/graphene square.
- Carefully remove the coverslip and ensure that the square adheres completely to the side of the coverslip upon removal.
- Transfer MMA/graphene squares to a clean 50 mL beaker filled with molecular grade water for 20 minutes.
- Gently plunge the coverslip with an MMA/graphene square attached into the water vertically and ensure that the square dislodges from the coverslip upon interaction with the surface of the water.
- Repeat for all MMA/graphene squares.

#### 1.7 Adhere graphene to grids

- Using negative action tweezers, gently dip a grid vertically into the water with the carbon side facing a floating MMA/graphene square.
- Once in contact with the square, carefully remove the grid and ensure that the square adheres completely to the carbon side of the grid upon removal. **CRITICAL:** Minimize lateral movement of the grid when submerged in water to prevent damaging grid squares.
- Place grids on a clean coverslip MMA/graphene side up and air dry for 1 to 2 minutes.
- Transfer coverslip graphene/MMA coated grids to a hot plate set to 130°C using flat-tip tweezers.
- Cover with the top of a glass petri dish and incubate for 20 minutes.
- Remove from heat and cool to room temperature for 1 to 2 minutes. **CAUTION:** Exercise caution as the petri dish will be extremely hot.

#### 1.8 Dissolve MMA and clean grids

- Transfer entire coverslip of grids into a petri dish filled with 15 mL acetone.
- Incubate for 30 minutes.
- Using a clean glass serological pipette, transfer acetone to a waste container.
- Carefully add 15 mL fresh acetone back to the petri dish using a clean glass serological pipette.
- Incubate for 30 minutes.
- Repeat these steps for a total of 3 acetone washes.
- Following 3 acetone washes, remove acetone and replace with 15 mL of isopropanol using a clean glass serological pipette.
- Incubate for 20 minutes.
- Repeat three times for a total of 4 isopropanol washes.
- Carefully remove grids from isopropanol and air dry on a clean coverslip using tweezers. **NOTE:** Residual isopropanol can make it difficult to release grids from tweezers. Making contact between the grid and the clean coverslip prior to releasing the grid from the tweezers can alleviate this problem.
- Evaporate residual organics by transferring the coverslip with graphene-coated grids on a hotplate set to 100°C for 10 minutes.
- Cool to room temperature and store under vacuum until use.

## PROTOCOL

### 2 UV/ozone treatment of graphene-coated grids

#### 2.1 Transfer grids

- Take graphene-coated grids out of vacuum storage and move grids into a temporary storage container (*e*.*g*. empty grid case, or a glass coverslip inside a petri dish). **CRITICAL:** Be sure to keep track of which face of the grid is coated with graphene.

#### 2.2 Prepare UV/ozone cleaner

- Slide open the illumination area of the UV/ozone cleaner and clean the surface with a Kimwipe.

#### 2.3 Insert grids

- Using reverse-action tweezers, gently place grids onto the illumination area. **NOTE:** Ensure the graphene-coated side is facing up.
- Slide the illumination area closed and turn the machine on.

#### 2.3 Treat grids

- Turn the time dial to the designated treatment time to begin UV/ozone treating the grids. This automatically begins the treatment. **NOTE:** Treatment duration is a tunable parameter, with excess treatment having the potential to damage the grid. Han *et al*. recommends 10 minutes of UV/ ozone treatment, and we have also found a 10-minute treatment to suffice. See Supplemental Protocol 1 for guidance on tuning this treatment time.

#### 2.5 Retrieve grids

- After the course of treatment has completed, turn the machine off and open the illumination area.
- Carefully retrieve the grids and place back them back into a temporary storage container, noting which side has been UV/ozone treated.

#### 2.6 Use grids

- Proceed directly to plunge-freezing a cryo-EM sample on the treated grids by, for example, following Vitrobot plunge-freezing steps as described in Koh *et al*^*37*^. **NOTE:** Atmospheric hydrocarbons can accumulate on the grid surface after UV/ ozone treatment and increase the hydrophobicity of the graphene surface^38,39^. To avoid this, we advise users to plunge the UV/ozone treated grids immediately. It may be advantageous to UV/ozone treat grids in batches, for example, UV/ ozone treating six and immediately plunging those six grids, then UV ozone treating a second batch of grids, followed by plunging the second batch.

## PROTOCOL

### 3 Diffraction analysis to assess graphene

#### 3.1 Tune the cryo-electron microscope

- Ensure that the microscope is well tuned with parallel illumination established.
- Correct defocus and set final defocus to -0.2 μm.

#### 3.2 Enter diffraction mode

- Insert the fluorescent screen.
- In the TEM User Interface (TUI) software insert the beam stop fully.
- On the instrument control panel select “Diffraction”. **NOTE:** Graphene spots should be clearly visible on the fluorescent screen. The diffraction beam may be off center. If this is the case, it should be shifted such that the highest intensity signal at the center of the diffraction pattern is blocked by the beam stop.

#### 3.3 Capture a diffraction image

- Within the Camera panel in TUI, insert CCD camera and lift the fluorescent screen.
- Set integration/acquisition time to 2 seconds.
- Press Acquire.

#### 3.4 Evaluate the diffraction image for quality

- Navigate to TEM Image Analysis software (TIA) to view the diffraction image. **NOTE:** The most easily observed diffraction spots generated by a graphene monolayer are six spots corresponding to a spatial frequency of 2.13 Å.
- Within TIA, use the measurement tool to estimate the distance from the center of the diffracted beam to one of these diffraction spots in reciprocal units. **NOTE:** 2.13Å≅0.47Å^-1^

## REPRESENTATIVE RESULTS

Successful fabrication of graphene-coated cryo-EM grids using the equipment (Figure 1) and protocol (Figure 2) outlined here will result in a monolayer of graphene covering the foil holes that can be confirmed by its characteristic diffraction pattern. To promote protein adsorption to the graphene surface, UV/ozone treatment can be used to render the surface hydrophilic by installing oxygen-containing functional groups. However, hydrocarbon contaminants in the air can adsorb onto the graphene surface as early as five minutes post UV/ ozone treatment and counteract this effect^38,39^. Importantly, both the duration of UV/ozone treatment and the time elapsed between treatment and plunging can affect sample quality. We demonstrate these effects using a simple method for assessing the hydrophilic character of the coated grid based on the surface contact angle (Figure 3; see Supplemental Protocol 1).

**Figure 1:**
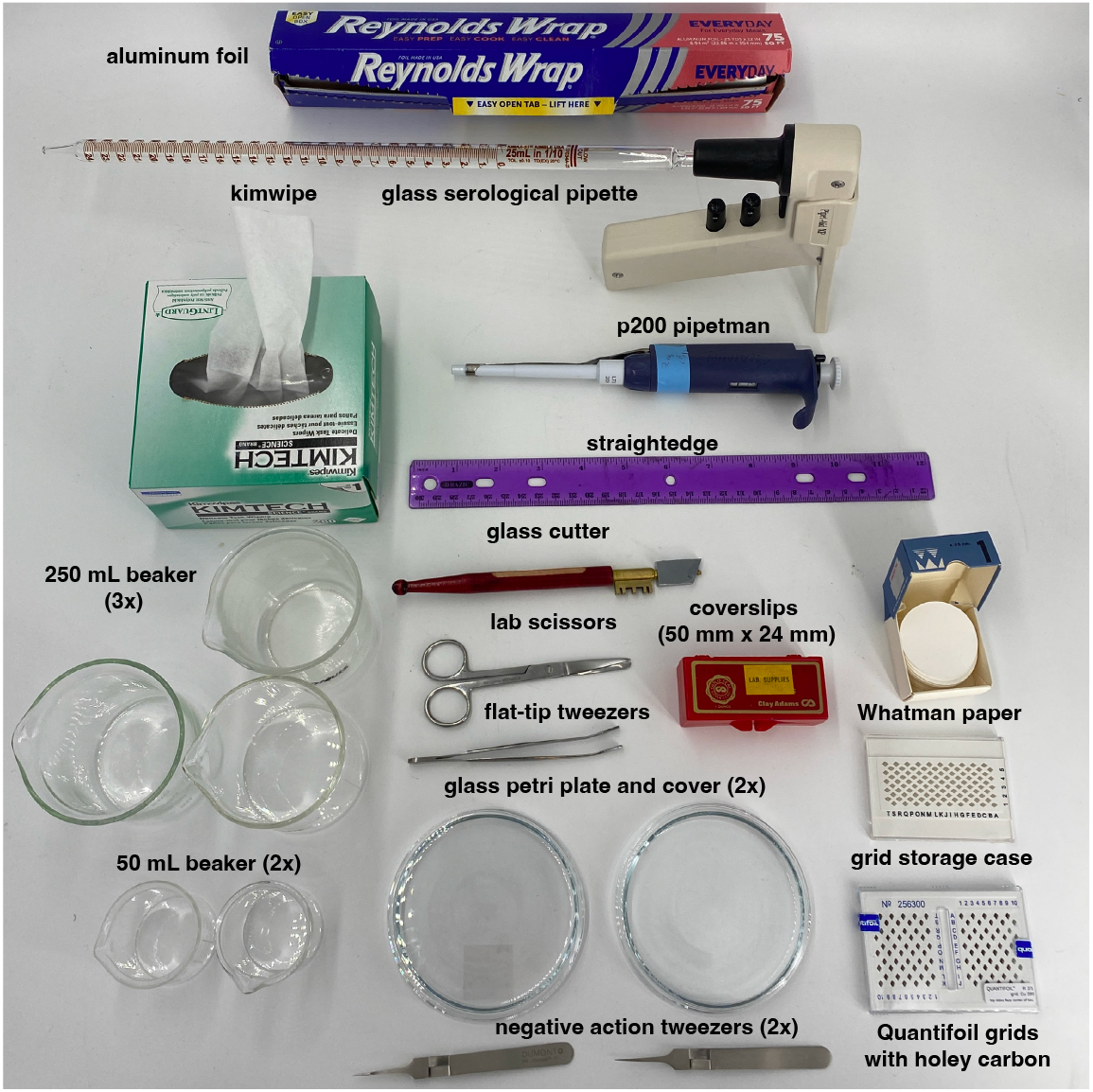
Required equipment. Lab equipment and tools necessary for the fabrication of graphene grids using the protocol detailed in this article. Items and their quantity are shown and labeled accordingly. Requisite reagents that are not shown include: CVD graphene, methyl-methacrylate EL-6 (MMA), ammonium persulfate (APS), acetone, isopropanol, ethanol, molecular grade water. Requisite instruments that are not shown include: spin coater, easiGlow discharger, hot plate, vacuum desiccator, and thermometer. All requisite items are detailed in the Table of Materials.

**Figure 2:**
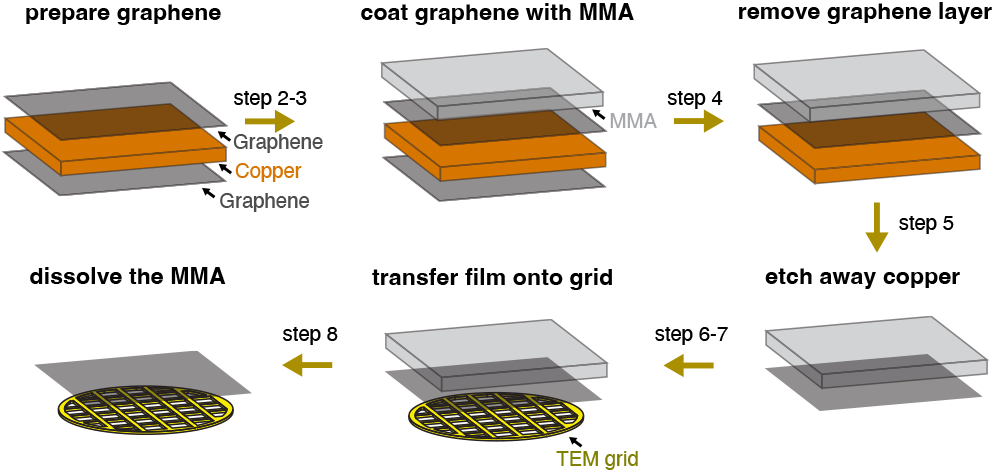
Schematic of the graphene grid fabrication process. Graphene is coated with a thin layer of methyl-methacrylate EL-6 (MMA) using a spin coater (steps 2-3). Graphene on the opposite side of the copper foil is removed via plasma etching (step 4). Ammonium persulfate (APS) is then used to etch away the copper (step 5). The MMA-graphene film is placed onto the grid surface (step 6-7). Lastly, MMA is dissolved during a series of washes with organic solvents (step 8). Steps indicated above arrows correspond to numbered steps described in protocol 1. This method is adapted from Han *et al*^30^.

**Figure 3:**
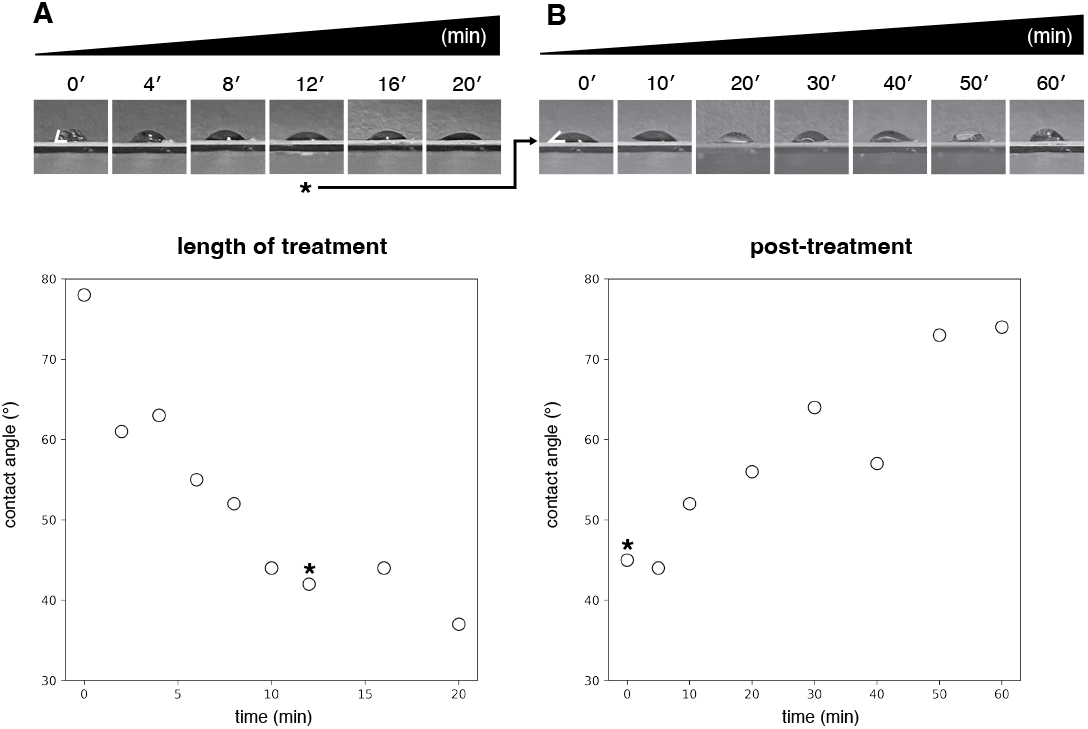
Assessment of grid surface hydrophilicity as a function of duration of UV/ozone treatment and time elapsed post-treatment. **A)** Measured contact angles plotted as a function of the duration of treatment. Decreased contact angles are consistent with increased hydrophilicity (untreated grid: 78°; 20 min treatement: 37°). Contact angles measured using ImageJ^43^. **B)** Measured contact angles plotted as a function of time, post treatment (0 min: 45°; 60 min: 74°). Grid measured in the post treatment time-course was UV/ozone treated for 12 minutes, as indicated by asterisk. Each post-treatment measurement was performed on the same grid, with the sample removed by wicking between measurements. Specific contact angles measured are expected to vary as a function of laboratory environmental conditions, and we recommend that users perform similar experiments in their laboratories to identify suitable conditions. See Supplemental Protocol 1 for more information.

To demonstrate the use of graphene supports in single particle cryo-EM, we applied a catalytically inactive RNA-guided DNA endonuclease *S. pyogenes* Cas9 (H10A; C80S; C574S; H840A)^40^ in complex with sgRNA and target DNA to graphene-coated grids, collected a cryo-EM dataset from these grids, and performed single particle analysis using the cryoSPARC software suite^7^. Graphene-coated grids consistently contained ∼300 particles per micrograph at 0.654 Å/pix magnification using a Titan Krios 300 keV microscope equipped with a Gatan BioQuantum-K3 detector (Figure 4A-E). An 8-hour data collection session with a +18° stage tilt yielded 2,963 movies and 324,439 particles in a final curated stack. Using these particles, we generated a 3D reconstruction which, upon refinement, yielded a density map with an estimated resolution of 2.7 Å and adequate angular sampling to avoid anisotropic artifacts (Figure 4F, 4H, 4I). An atomic model (PDB 6o0z)^41^ was docked into this map, and refined using ISOLDE^42^. Residues R63-L82 of this fitted atomic model are displayed with the refined cryo-EM density map, highlighting the resolved side-chain density (Figure 4F). When comparing the same sample and concentration (250 ng/μL) applied to identical grids that lacked graphene, no particles were observed (Figure 4G). This observation highlights the efficacy of the graphene support in enabling the visualization of particles from low-concentration samples.

**Figure 4:**
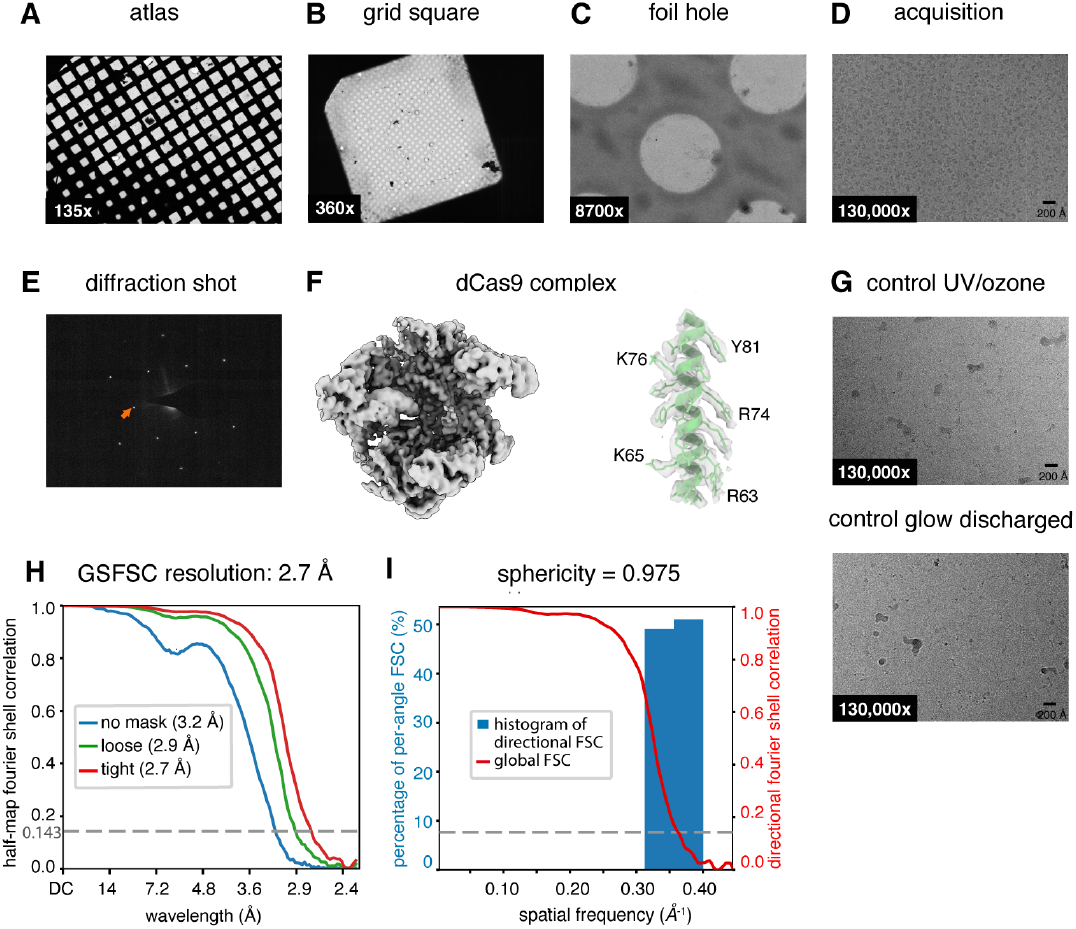
Representative images of graphene-coated grid and cryo-EM reconstruction of a Cas9 complex. **A-C)** Representative atlas, grid square, and foil hole images of graphene-coated holey carbon Quantifoil grids taken on a Titan Krios 300 keV microscope equipped with a Gatan BioQuantum-K3 detector. **D)** Cryo-EM micrograph of *S. pyogenes* dCas9 in complex with sgRNA and target DNA (conc. 250 ng/μL) on graphene-coated holey carbon grid. **E)** Diffraction image from grid imaged in panels A-D. Orange arrow indicates a position corresponding to a spatial frequency of 2.13 Å. **F)** Cryo-EM density map from 3D reconstruction of the dCas9 complex (left). Residues R63-L82 from a fitted model are depicted within the semi-transparent cryo-EM density, with a subset of visible side-chains labeled (right). **G)** An identical sample to that in panel D was applied to UV/ozone treated (top) and glow discharged (bottom) holey carbon grids without graphene. Cryo-EM micrographs displayed are representative of each grid and show no particles. **H)** Fourier Shell Correlation (FSC) curves of the unmasked, loosely, and tightly masked maps computed in CryoSPARC. **I)** Histogram and directional FSC plot computed in CryoSPARC based on the 3DFSC method^23^. See Supplemental Figure 2 and Supplemental Protocol 2 for more information.

## DISCUSSION

Cryo-EM sample preparation involves a host of technical challenges, with most workflows requiring researchers to manually manipulate fragile grids with extreme care to avoid damaging them. Additionally, the amenability of any sample to vitrification is unpredictable; particles often interact with the air-water-interface or with the solid support foil overlaying the grids, which can lead to particles adopting preferred orientations or failing to enter the imaging holes unless very high protein concentrations are applied^24^. Overlaying holey cryo-EM grids with a continuous monolayer of graphene has shown tremendous promise in improving particle distributions on micrographs, increasing particle numbers at low concentrations and reducing preferred orientations driven by interactions at the air-water-interface^30^.

A limitation of existing graphene coating protocols for cryo-EM grids is the extensive manual manipulations required for the coating process, which can compromise quality and increase grid-to-grid variability. In this work we describe slight modifications to an existing protocol for coating cryo-EM grids with a monolayer of graphene^30^. These modifications emphasize caution in grid handling and reduce manual manipulations of the grids during graphene application, annealing, and washing, thereby aiming to increase the yield of intact, high quality, graphene-coated grids.

Graphene-coated grids typically require UV/ozone treatment to render the surface hydrophilic for sample application. The duration of UV/ozone treatment and the time elapsed after treatment and prior to plunging can impact grid hydrophilicity and ultimately sample quality. In addition to the grid fabrication protocol, we describe a technique for assessing grid hydrophilicity following UV/ozone treatment. In the procedure, the surface contact angle of an applied sample is used as an indicator of the hydrophilic character of the coated grid^20,44^. Designs are provided to inexpensively 3D-print a custom grid imaging mount that utilizes a simple cell phone camera to estimate the surface contact angle.

Finally, we describe results obtained by employing this protocol to determine the 2.7 Å cryo-EM structure of the catalytically inactive RNA-guided DNA endonuclease, *S. pyogenes* Cas9 in complex with sgRNA and target DNA^40^. In the absence of graphene, no particles were observed in foil holes at the complex concentrations used (250 ng/μL). In contrast, graphene-coated grids bore particles at high density, enabling facile 3D-reconstruction of a high-resolution map from 2,961 micrographs. Taken together, these data highlight the value of applying graphene monolayers to cryo-EM grids for single particle analysis.

## Supporting information

3D Printing Files

## ADDITIONAL MATERIALS

Stereolithography CAD files in the STL format are provided to facilitate 3D printing of the tabletop imaging mount (slide_mount_v1.stl) and camera stand (camera_stand_v1.stl). Devices shown in this protocol were printed using a Prusa i3 MK3s printer.

## ACKNOWLEDGMENTS

Specimens were prepared and imaged at the Cryo-EM Facility in MIT.nano on a Titan Krios microscope, and on a Talos Arctica microscope, which was a gift from the Arnold and Mabel Beckman Foundation. Contact angle imaging devices were printed at the MIT Metropolis Maker Space. We thank the laboratories of Nieng Yan and Yimo Han, and staff at MIT.nano for their support throughout the adoption of this method. In particular, we extend our thanks to Drs. Guanhui Gao and Sarah Stirling for their insightful discussions and feedback. This work was supported by NIH grants R01-GM144542, 5T32-GM007287, and NSF-CAREER grant 2046778. Research in the Davis lab is supported by the Alfred P. Sloan Foundation, the James H. Ferry Fund, the MIT J-Clinic, and the Whitehead Family.

## TABLE OF MATERIALS

| Name of Material/Equipment | Company | Catalog Number | Comment |
| --- | --- | --- | --- |
| Acetone | Fisher | A949-4 |  |
| Aluminum foil | Fisher | 15-078-292 |  |
| Ammonium persulfate | Fisher | (117874 |  |
| Coverslips 50 mm x 24 mm | Mattek | PCS-1.5-5024 |  |
| CVD graphene | Graphene Supermarket | CVD-Cu-2x2 |  |
| easiGlow discharger | Ted-Pella | 91000S |  |
| Ethanol | Millipore-Sigma | 1.11727 |  |
| Glass cutter | Grainger | 21UE26 |  |
| Glass serological pipette | Fisher | 13-676-34D |  |
| Grid Storage Case | EMS | 71146-02 |  |
| Flat-tip tweezers | Fisher | 50-239-60 |  |
| Hot plate | Fisher | 07-770-108 |  |
| Isopropanol | Sigma | W292907 |  |
| Kimwipe | Fisher | 06-666 |  |
| Glass petri dish and cover | VWR | 75845-544 |  |
| Lab scissors | Fisher | 13-806-2 |  |
| Methyl-Methacrylate EL-6 (MMA) | Kayaku | M310006 0500L1GL |  |
| Molecular grade water | Corning | 46-000-CM |  |
| Negative action tweezers (2x) | Fisher | 50-242-78 |  |
| Parafilm | Fisher | 13-374-12 |  |
| P20 pipette | Rainin | 17014392 |  |
| P200 pipette | Rainin | 17008652 |  |
| Pipette tips | Rainin | 30389291 |  |
| Quantifoil grids with holey carbon | EMS | Q2100CR1 |  |
| Spin coater | SetCas | KW-4A | with chuck SCA-19-23 |
| Straightedge | ULINE | H-6560 |  |
| Thermometer | Grainger | 3LRD1 |  |
| UV/Ozone cleaner | BioForce | SKU: PC440 |  |
| Whatman paper | VWR | 28297-216 |  |
| Vacuum desiccator | Thomas Scientific | 1159X11 |  |
| 50 mL beaker (2x) | Corning | 1000-50 |  |
| 250 mL beaker (3x) | Fisher | 02-555-25B |  |

## FIGURES

**Supplementary Figure 1:**
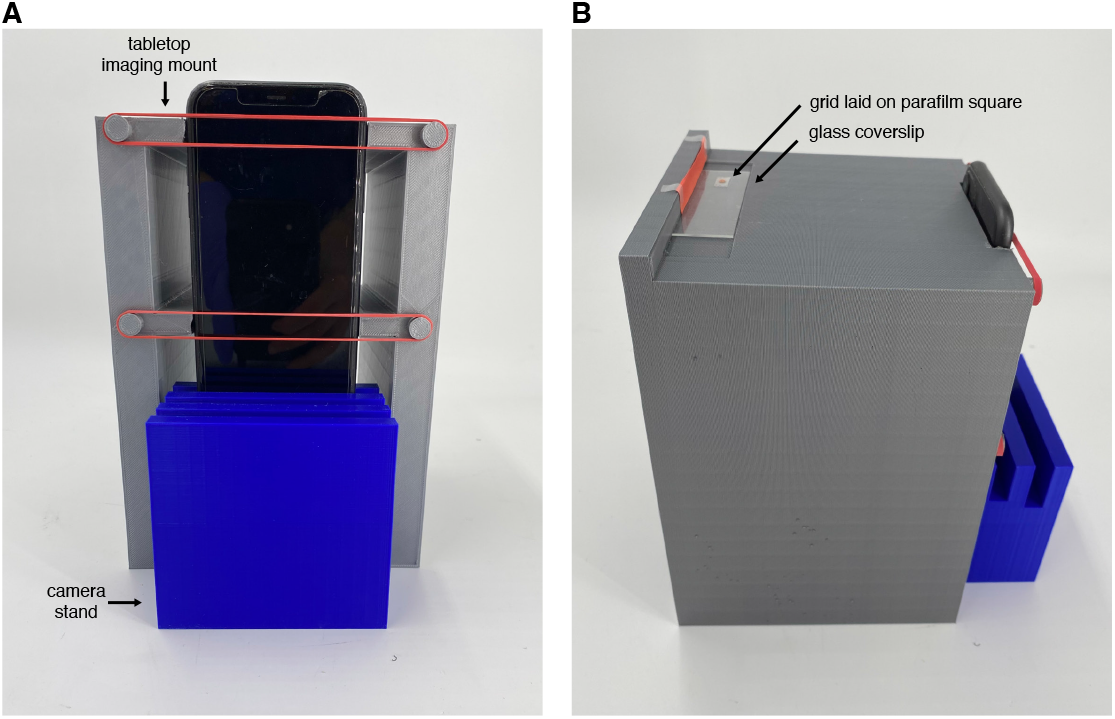
Contact angle imaging stand. **(A)** 3D printed camera stand and tabletop imaging mount to secure camera in a position that aligns the camera in-plane with the coverslip. **(B)** Grid is placed on the coverslip on top of a 1 cm x 1 cm square piece of parafilm. Depicted imaging mount was 3D-printed using the provided .stl files, and uses an iPhone 11, but can be readily modified to accommodate other devices.

**Supplementary Figure 2:**
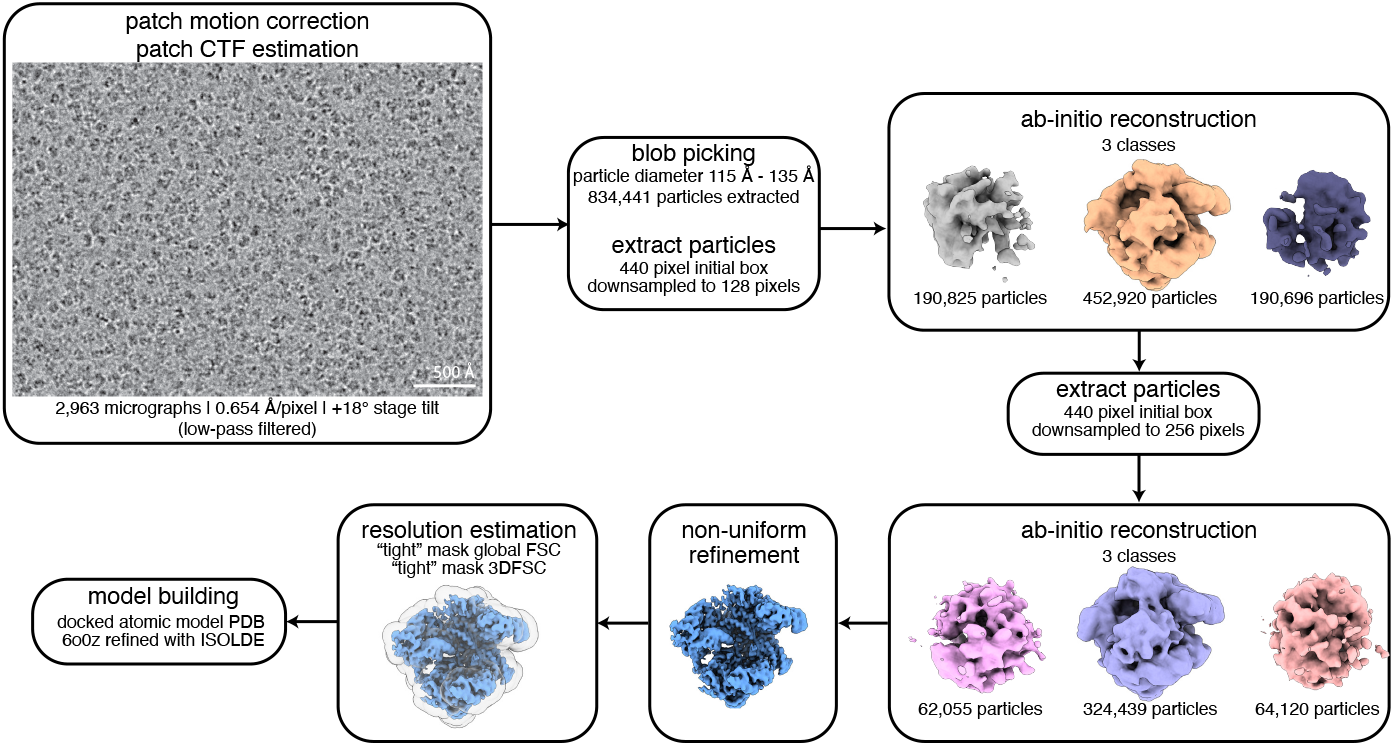
Image processing workflow. CryoSPARC (v4.2.1) processing workflow for dCas9 complex. Job names, job details, and non-default parameters (italicized) are indicated.

## SUPPLENTARY PROTOCOL

**<u>Protocol S1. Assessing grid hydrophilicity</u>**

**S1.1** Prepare imaging setup.

- Locate a phone stand, tabletop imaging surface, phone with camera, glass coverslip, and parafilm. **NOTE:** Images shown in Figure 3 were obtained using a phone stand and tabletop imaging surface that were 3D printed using the provided .stl files, resulting in more easily reproducible and quantifiable contact angle measurements (**Figure S1A**).
- Lay a glass coverslip onto the flat surface.
- Cut a 1 cm by 1 cm square of parafilm and place it onto the glass coverslip.
- Place phone on the phone stand, orienting the phone such that the camera is inplane with the glass coverslip. Secure phone in this position using rubber bands (**Figure S1B)**.

**S1.2** Image grid.

- Take a sample photo to verify that the camera is aligned with the imaging surface.
- Directly after UV/ozone treatment of graphene-coated grids (see Protocol 2), place a single grid onto the square of parafilm on the glass coverslip. **CRITICAL:** Ensure the graphene side is facing upwards.
- Add a 2 μL water droplet onto the center of the grid surface with a pipette and immediately take a photo.

**S1.3** Test treatment conditions.

- Repeat steps 1.5-1.6 after: *i*) desired intervals of UV/ozone treatment to determine a sufficient length of treatment; or *ii*) desired time intervals post UV/ozone treatment to measure how long after treatment the grid surface maintains its hydrophilic character.

**S1.4** Quantify hydrophilicity.

- Calculate contact angles from photos by importing them into ImageJ^43^ and using the Contact Angle plugin.

## SUPPLEMENTARY PROTOCOL

**<u>Protocol S2. Single particle analysis of the dCas9 complex dataset</u>**.

**NOTE:** All image processing described in this protocol was performed using cryoSPARC version 4.2.1.

**S2.1** Preprocess movies using the “Patch Motion Correction” and “Patch CTF Estimation” jobs.

**S2.2** Perform particle picking using the “Blob picker” job, using a spherical blob ranging in diameter from 115 Å to 135 Å.

**S2.3** Extract particles using the “Extract from micrographs” job, using normalized correlation coefficient (NCC) and power thresholds resulting in approximately 200-300 particles per micrograph.

**NOTE:** Appropriate thresholds and resulting particle count may vary, and users should inspect pick quality across a range of micrographs to identify suitable conditions.

**S2.4** Perform multiclass initial reconstructions using the “Ab initio reconstruction” job, requiring three classes. Two of the three classes will likely contain non-Cas9 particles, including surface contaminants. Select the class resembling dCas9 for further processing. Additional rounds of multiclass “Ab initio reconstruction” or “Heterogeneous refinement” may be applied to further refine the particle stack.

**S2.5** Perform 3D refinement using the “Non-uniform Refinement” job selecting default parameters.

**S2.6** Estimate resolution of the reconstruction using the “Validation (FSC)”, and “ThreeDFSC” jobs, employing the maps and mask from the final 3D refinement.

